# Longitudinal fluctuations in protein concentrations and higher-order structures in the plasma proteome of kidney failure patients subjected to a kidney transplant

**DOI:** 10.1101/2024.01.31.578168

**Authors:** Sofia Kalaidopoulou Nteak, Franziska Völlmy, Marie V. Lukassen, Henk van den Toorn, Maurits A. den Boer, Albert Bondt, Sjors P.A. van der Lans, Pieter-Jan Haas, Arjan D. van Zuilen, Suzan H. M. Rooijakkers, Albert J.R. Heck

## Abstract

Using proteomics and complexome profiling we evaluated over a period of a year longitudinal variations in the plasma proteome of kidney failure patients, prior to and after a kidney transplantation, comparing this data with two healthy controls. The post-transplant period was complicated by numerous bacterial infections, resulting in dramatic changes in the plasma proteome, mostly related to an acute phase condition. As positive acute phase proteins, being elevated upon inflammation, we observed the well-described C-reactive protein (CRP) and Serum Amyloid A (SAA1 and SAA2), but our analyses added to that Fibrinogen (FGA, FGB and FGG), Haptoglobin (HP), Leucine-rich alpha-2-glycoprotein (LRG1), Lipopolysaccharide-binding protein (LBP), Alpha-1-antitrypsin (SERPINA1), Alpha-1-antichymotrypsin (SERPINA3), Protein S100 (S100A8, S100A9), Complement protein C4, C4b-binding protein alpha chain (C4BPA), Complement factor B (CFB) and Monocyte differentiation antigen CD14. As negative acute phase proteins, being downregulated upon inflammation, we identified the well-documented Serotransferrin (TF) and Transthyretin (TTR), but add on to that Kallistatin (SERPINA4), Heparin cofactor 2 (SERPIND1), Inter-alpha-trypsin inhibitor heavy chain H1 and H2 (ITIH1, ITIH2). For a patient with the most severe acute phase response, we furthermore performed plasma complexome profiling by SEC-LC-MS on all longitudinal samples. We observe that several plasma proteins displaying alike concentration patterns, co-elute and putatively form macromolecular complexes. These include a) FGA, FGB and FGG (as expected, b) ITIH1 and ITIH2, c) HP together with Hemoglobin (HB), d) the small acute phase biomarker proteins SAA1 and SAA2 with the Apolipoproteins A-I, A-II, A-IV (APOA1, APOA2, APOA4). By complexome profiling we expose how SAA1 and SAA2 become incorporated into high-density lipid particles, thereby replacing partly APOA1 and APOA4. Overall, our data highlight that the combination of in-depth longitudinal plasma proteome and complexome profiling can shed further light on the correlated variations in the abundance of several plasma proteins upon inflammatory events.

## Introduction

Plasma proteomics has matured substantially and can nowadays be used to assess changes in the plasma proteome of hundreds of samples taken from dozens of patients to monitor changes within and between donors under different physiological conditions^1–4^. In plasma proteomics the abundances of the proteins are mostly assessed over large cohorts, albeit taking often just a single (or a few) sampling timepoint for each donor. Moreover, the abundance of each protein is measured as being a discrete entity, ignoring that many plasma proteins are organized as multi-component systems, with IgM and haemoglobin (HB) being illustrative examples^5, 6^. Here we report an alternative in-depth plasma proteomics study in which we focus on these latter aspects. We study the changes in the plasma proteome of two healthy controls and two chronic kidney disease (CKD) patients, who underwent a kidney transplant and consequently, suffered from bacterial infections.

Chronic kidney disease (CKD) is a severe disorder, affecting almost 600 million people worldwide^7^. CKD has been described as a major contributor of mortality globally^8^ and it is thus crucial to improve its diagnosis and treatment methods. Upon progression, CKD requires renal replacement therapy via either dialysis treatment or kidney transplantation. The latter is usually performed in patients that have already been on dialysis. This operation is risky, as several immune responses can take place in response to the foreign organ, alike to adaptive responses during microbial infections^9^.

CKD is associated with disturbances in markers of inflammation, lipid metabolism other metabolic pathways and the accumulation of water-soluble waste products which normally are excreted via urine. Moreover, the presence of proteinuria in some CKD patients has impact on their plasma proteome, by the loss of specific proteins^10^. Also, dialysis and other issues related to the care of patients with chronic kidney disease (CKD), such as drug-treatments, may impact their plasma proteome.

After kidney transplantation, various features which may impact the patients’ plasma proteome come together. The transplantation process itself with the introduction of an organ from a donor can induce changes in the plasma proteome, both through ischemic reperfusion injury, allorecognition and even a possible rejection. Furthermore, the drugs used to protect the kidney transplant recipient from rejection or opportunistic infections themselves could influence the patients’ plasma proteome. Finally, patients using immunosuppressive therapy are at risk for several infectious complications, both of viral and bacterial origin. In the case of kidney transplantation, urinary tract infections occur frequently. This infectious disease typically results in an inflammatory response.

A key component of the inflammatory response upon bacterial infection, is the acute phase response (APR). The APR is characterized by at least 25% increase or decrease of the positive or negative acute phase proteins^11^. Some of these Acute Phase Proteins (APPs), have already been identified as biomarkers, like CRP and SAA, whose concentrations can increase 1000-fold during an APR. APR has already been studied extensively for decades, with major focus on alterations in the concentrations of specific proteins, measuring one at the time. Here we aimed to longitudinally monitor the APR in two chronic kidney disease patients who suffered from bacterial infection in the months after the kidney transplantation. By combining quantitative proteomics to measure simultaneously protein concentrations of hundreds of plasma proteins with complexome profiling^12^, we aim to explain certain mechanisms behind protein variations during the APR, focusing on how protein associations correlate with changing protein abundances.

To achieve the aforementioned goal, we monitored the proteome profile of two patients right before and after kidney transplantation, through a time span of a year after surgery. For reference we also took longitudinal samples from two healthy controls along. Using the robust and sensitive data-independent acquisition (DIA) approach we were able to identify specific concentration patterns of both abundant and less abundant proteins in the patients before transplantation, during infections and between the healthy and the diseased donors. The quantitative longitudinal plasma proteomics data allowed the classification of many known positive and negative APPs and revealed some putative new APR proteins. Furthermore, we used size exclusion chromatography (SEC) to separate native plasma proteins and protein complexes. We correlated plasma protein complexes with changes in abundance, which, among others, clearly revealed the co-elution of SAA with the HDL particles, thereby replacing specific Apolipoproteins upon the APR.

## Materials and methods

### Human plasma samples

The plasma samples for the two healthy controls were obtained from PrecisionMed, Carlsbad, California. For both these donors, plasma was obtained from 3 consecutive samplings, each time a month apart. We designate these donors as control C1 and C2, and the sampling times T0-T2. These plasma samples therefore cover in total a period of two months. These samples were included to provide baseline levels of plasma protein concentrations, and to assess for biological variability between these healthy donors and over time. The two kidney transplant recipients, here after termed P1 and P2, both received a kidney transplant, shortly after the sampling of the first blood sample, here termed T0. After transplantation, these two patients were closely monitored for infections, and their blood (and urine) was routinely sampled over a period of close to a year to monitor viral reactivations. For P1 we obtained nine post-transplantation EDTA plasma samples (named T1–T9), and for P2 four (named T1-T4).

### Plasma sample preparation for DIA LC-MSMS

As adapted by Völlmy et al^4^, 24 μL of a detergent-based buffer (1% sodium deoxycholate (SDC), 10 mM tris (2-carboxyethyl) phosphine (TCEP), 10 mM Tris, and 40 mM chloroacetamide) with Complete mini EDTA-free protease inhibitor cocktail (Roche) was added to 1 μL plasma and boiled for 5 min at 95°C to enhance protein denaturation. 50 mM ammonium bicarbonate was added, and digestion was allowed to proceed overnight at 37°C using trypsin (Promega) and LysC (Wako) at 1:50 and 1:75 enzyme:substrate ratios, respectively. The digestion was quenched with 10% formic acid (FA) and the resulting peptides were cleaned-up in an automated fashion using the AssayMap Bravo platform (Agilent Technologies) with a corresponding AssayMap C18 reverse-phase column. The eluate was dried and resolubilized in 1% FA to achieve a concentration of 1 µg/μL, of which 1 μL was injected.

### LC-MS/MS data-independent acquisition

All spectra were acquired on an Orbitrap Exploris 480 mass spectrometer (Thermo Fisher Scientific) operated in data-independent mode (DIA) coupled to an Ultimate3000 liquid chromatography system (Thermo Fisher Scientific) and separated on a 50-cm reversed phase column packed in-house (Poroshell EC-C18, 2.7 μm, 50 cm × 75 μm; Agilent Technologies). Proteome samples were eluted over a linear gradient of a dual-buffer setup with buffer A (0.1% FA) and buffer B (80%ACN, 0.1%FA) ranging from 9 to 44% B over 65 min, 44–99% B for 3 min, and maintained at 95% B for the final 5 min with a flow rate of 300 nl/min. DIA runs consisted of a MS1 scan at 60,000 resolution at m/z 200 followed by 30 sequential quadrupole isolation windows of 20 m/z for HCD MS/MS with detection of fragment ions in the Orbitrap (OT) at 30,000 resolution at *m/z* 200. The *m/z* range covered was 400–1,000 and the Automatic Gain Control was set to 100% for MS and 1000% for MS/MS. The injection time was set to “custom” for MS and “auto” for MS/MS scans.

### Raw data processing

Spectra were extracted from the DIA data using DIA-NN (version 1.8) without a spectral library and with “Deep learning” option enabled^13^. The enzyme for digestion was set to trypsin and one missed cleavage was tolerated. Cysteine Carbamidomethylation and Methionine oxidation were allowed. The precursor false discovery rate threshold was set to 1%. Protein grouping was done by protein names and cross-run normalization was RT-dependent. The MS1 mass accuracy was 4.7 ppm based on the first run of the experiment and for MS2 the optimised mass accuracy was set at 14.8 ppm. All other settings were kept at the default values and can be found in the log files on the PRIDE Submission, which can be accessed through the Data Availability section. The gene-centric report from DIA-NN was used for downstream analysis, and quantification was based on unique peptides. When injection replicates were available, the median of these values was used. The FT and T10 samples were excluded from the statistical analysis as they were not relevant for the current study. The Uniprot reviewed human protein database was used, with ∼ 20300 entries (Release number 2018_05). An additional sequence of a dominant IGHG1 clone for P1 patient was added in the search, previously described by Peng et al^14^. All downstream analyses were carried out in R15.

### Plasma sample fractionation by Size Exclusion Chromatography (SEC)

18μL of plasma from every timepoint (T0, T1, T2, T3, T4) of patient P2 and plasma from T1 of the C1 healthy individual, were fractionated offline on an Agilent 1290 Infinity HPLC System using phosphate-buffered saline (PBS) as the mobile phase. The buffer was filtered using a 0.22-μm disposable membrane cartridge (Millipore). The fractions were eluted offline over a 60min gradient through a dual column set-up of a YarraTM 3 μm SEC-4000 (300 x 7.8 mm; Phenomenex) and a YarraTM 3 μm SEC-3000 (300 x 7.8 mm; Phenomenex), with a isocratic flow rate of 0.5 ml/min, as previous described by Tamara et al^6^. 96 fractions were collected in total over a 30 min collection time. Blue Dextran from the Gel Filtration Molecular Weight Markers Kit for Molecular Weights 29,000–700,000 Da (Sigma Aldrich) was used as a void volume marker and the six proteins (Albumin, Carbonic Anhydrase, Alcohol Dehydrogenase, β-Amylase, Apoferritin, Thyroglobulin) as calibration standards. The SEC-chromatograms were measured using absorption at 280 nm.

### LC-MS/MS DIA for fractionated SEC samples

The fractions with molecular weight between ∼ 1 kDa and ∼ 2 MDa were selected for analysis from all timepoints of all samples, cumulating in to 66 fractions per timepoint. The fractions were analysed using the same DIA method as described above on an Orbitrap 480 spectrometer (Thermo Scientific), coupled to an Evosep One liquid chromatography system, using the Endurance column (EV1106) at 30SPD method (44min gradient). The samples were loaded on Evotips C18 disposable trap columns (EV2018). The DIA method consisted of MS1 scans at 60,000 resolution and scan range between 375-1600, followed by 50 scan windows with 12 *m/z* and 1 window overlap in the HCD MS/MS. The resolution on the OT was set to 15,000 at 200 *m/z* with precursor mass range of 400-1000 *m/z*.

### Raw data processing and statistical analysis of data from SEC fractionated samples

All fractions of all time points within a sample were analysed in one run to avoid batch effects, using DIA-NN’s software (version 1.8.1)^13^. The software settings were the same as for the unfractionated samples, besides disabling all normalization methods. The MS1 mass accuracy was at set automatically to 1.5 ppm whereas the MS2 optimised mass accuracy was 1.6 ppm. Further details on DIA-NN’s runs can be found in the log files. The *evidence* output file of DIA-NN was used for the analyses, which was filtered at 1% Q-value and Lib PG Q-Value. The *PG Quantity* was used to calculate the abundance of the protein groups and the Precursor Intensity for the peptides. The Uniprot human protein database was used, with ∼ 103000 entries (Release number 2023_06). Certain fractions were removed from the subsequent analysis from all timepoints because of the pure quality of the chromatograms and/ or LC-MS runs likely due to errors in sample preparation (i.e., fractions 18, 21, 22, 31, 32, 50). Furthermore, keratins and variable regions covering the highly variable CDRs of immunoglobulins were removed.

## Results

### Characteristics of the two healthy controls and the two chronic kidney disease patients

After the kidney transplant both patients suffered from multiple bacterial infections, as evidenced by positive bacterial cultures when samples were extracted either from blood or urine. For P1, testing revealed that this was an invasive infection with *K. pneumoniae,* and also recurrent infections by a colonising strain. P2 had an infection with *K. pneumoniae* and a subsequent infection with *E. coli*. The characteristics of the timelines of these two patients are indicated in **Figure 1**, whereby the timing of the longitudinal plasma sampling is indicated, by T0-T9 and T0-T4, respectively for P1 and P2. For comparison the timepoints of sampling from the two healthy donors are also indicated in **Figure 1**.

**Figure 1.**
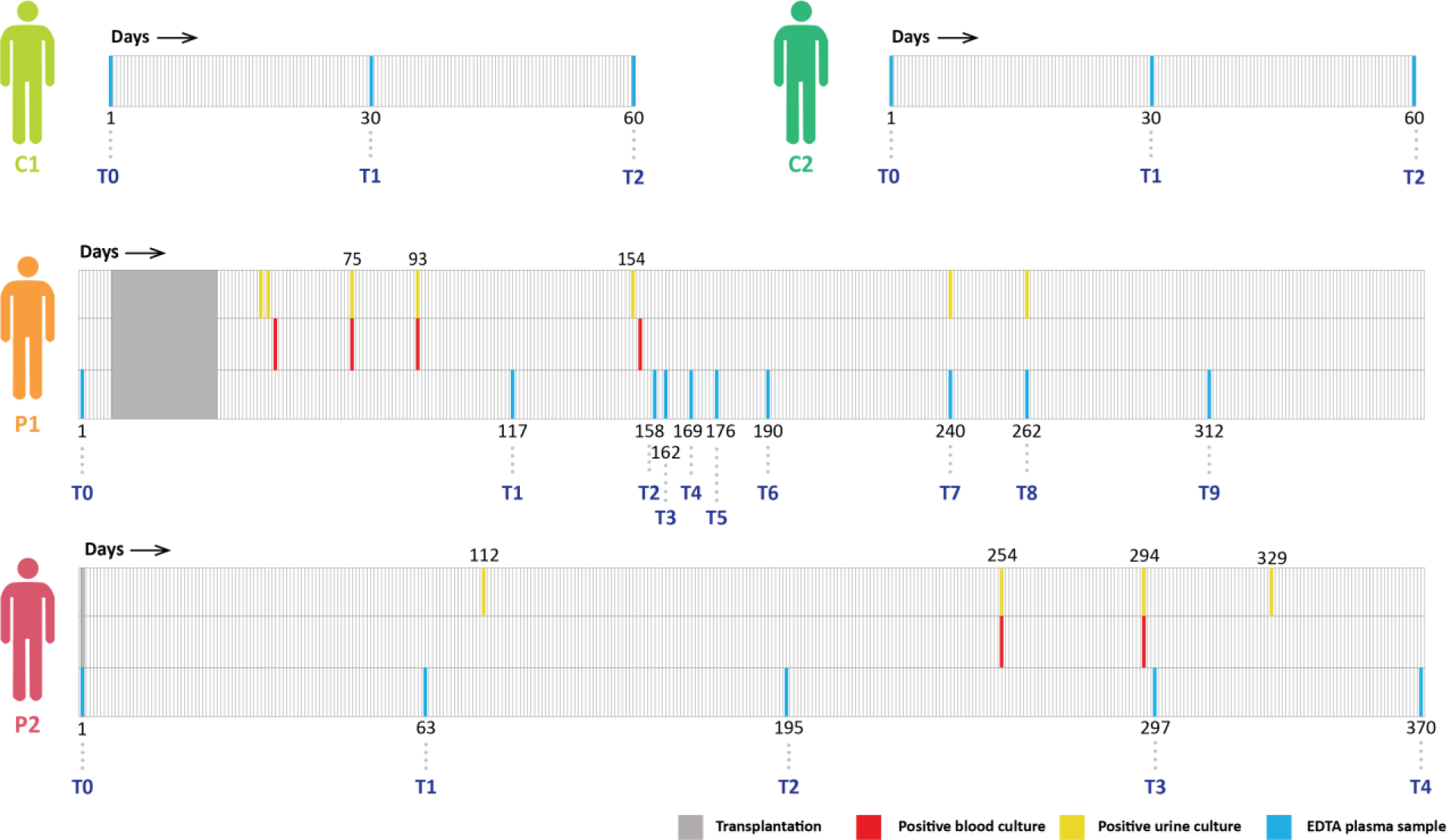
Timeline of drawing plasma samples and bacterial cultures from the kidney patients (P1, P2) and healthy control donors (C1, C2). The blue lines indicate the instants of blood sampling as used for the proteomics profiling, the red show the positive blood bacterial culture and the yellow the positive urine bacterial culture. All samples were taken in the span of about a year, with the number below the timelines depicting the number of days after the initial sampling, starting from day 1. The grey areas in the two patients indicate the time of the transplantation. For both patients the T0 timepoint was taken before transplantation, either the same day (P2) or a few days before (P1). For the healthy individuals (C1, C2) the exact day of the blood collection is unknown but the timepoints are about two consecutive months apart.

### Plasma Proteomics; determining concentrations of a few hundred of proteins over 21 plasma samples

In total we analysed 21 plasma samples, 3 from C1, 3 from C2, 10 from P1 and 5 from P2. We used a robust data-independent acquisition (DIA) approach similar to described earlier by Völlmy et al. ^4^ to profile the protein levels in plasma of the healthy donors and the two patients, as this approach largely circumvents the semi-stochastic sampling bias specific to data-dependent shotgun proteomics, and benefits from high reproducibility. In total we were able to quantify about 480 proteins across all 21 plasma samples, although this number still contains quite a few variable immunoglobulin protein fragments and protein isoforms (Supplementary Table 1). In plasma proteomics it has been well established that the total intensity of a protein in label free quantification (i.e., LFQ- or IBAQ-values) can be used as a proxy for protein concentrations. To better relate the abundance of plasma proteins to clinical data, we converted the median log label-free quantified values per protein from our LC-MS experiments into plasma protein concentrations. For this conversion, as described in more detail previously^4^, we performed a linear regression with 22 known reported average values of proteins in plasma (A2M, B2M, C1R, C2, C6, C9, CFP, CP, F10, F12, F2, F7, F8, F9, HP, KLKB1, MB, MBL2, SERPINA1, TFRC, TTR, VWF) ^16^. Supplementary Table 1 provides for all detected 480 proteins the determined concentrations over all time points both for the two healthy donors and the two patients. To provide an indication of the achievable dynamic range, the measured concentrations range from ∼ 2000 mg/dL for albumin to about 0.02 mg/dL for S100A8/S100A9.

### Global features observed in the plasma proteomes

To obtain a first global overview of the plasma-proteome profiles, we organized all data in **Figure 2**, whereby in **Figure 2a** we display a heatmap representation of the concentrations of the 200 unique and most abundant plasma proteins. Although this heatmap is not easy to interpret in detail, some global features become already directly evident. For instance, the plasma proteomes of the two control patients are quite stable over time (from T0 to T2), and even between C1 and C2 there is seemingly little variability. This is evidence that our quantitative plasma proteome data is quite accurate and reproducible and that the two healthy donors can genuinely be regarded as proper controls.

**Figure 2.**
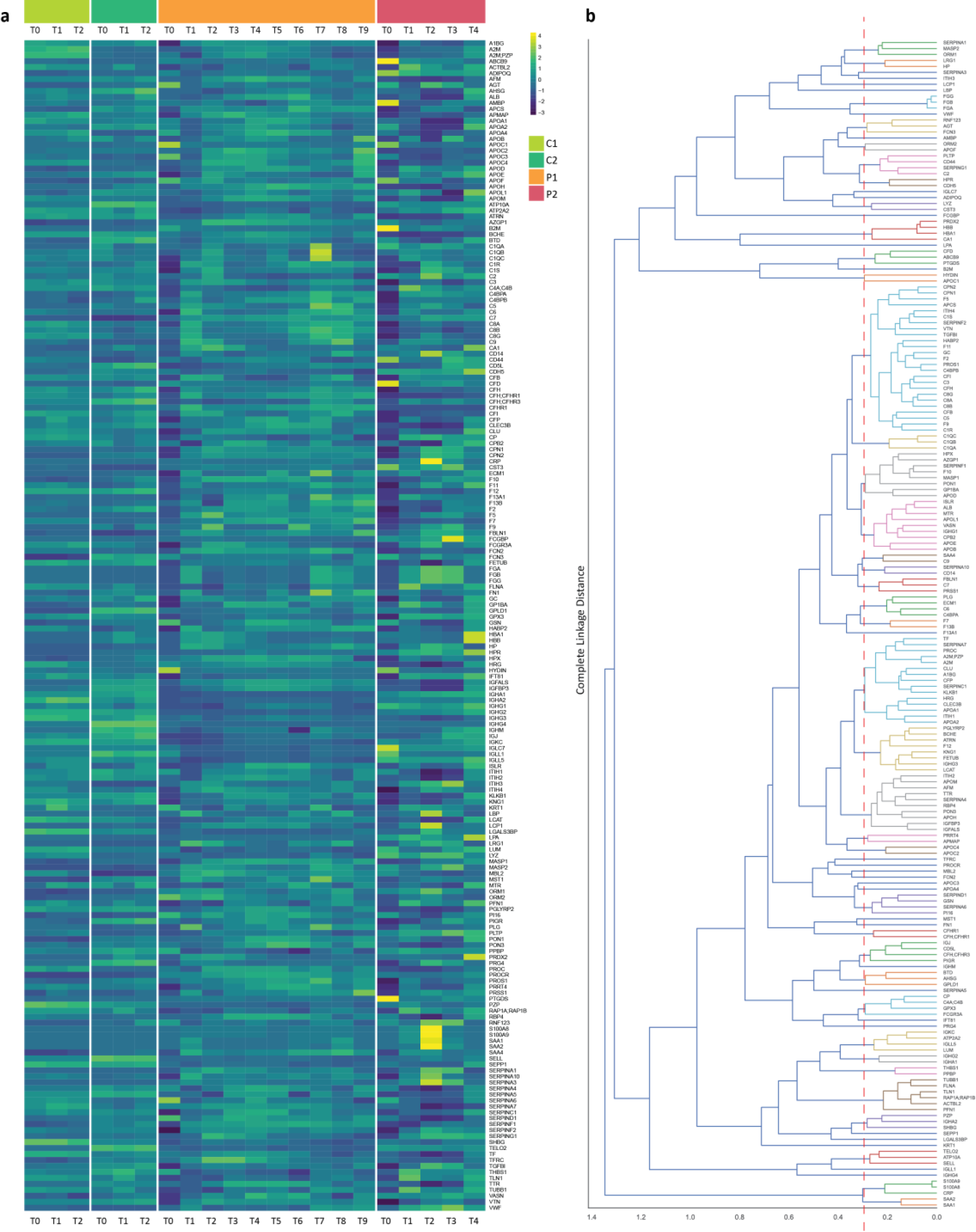
Visualization of variations in protein concentration of the 200 most abundant proteins in plasma. a. Heatmap with the abundance of proteins in each timepoint of each sample. Each individual is represented by a different colour on the top of the figure. The more yellow, the higher the concentration and the darker the blue the lower the concentration. Overall, the controls (C1,C2) show more alike profiles across the timepoints, whereas the patients (P1, P2) show substantial more variations. Moreover, timepoints T0 in the P1 and P2 show alike outlier profiles as well as T2 in P2. **b.** Dendogram that clusters the plasma proteins based on similarities in their plasma concentrations. The lower the complete distance value the more (cor)related the proteins likely are. Proteins with less than 0.3 distance are separated with a red dashed line. For example, the three chains of fibrinogen (light blue colour) are clustered tightly together within less than 0.1 distance. In the same way C1QA, C1QB and C1QC are clustered tight together (light yellow).

In contrast, the two patients show striking variation over time. The highest deviations were observed for the two patients’ samples taken before the surgery (P1_T0 and P2_T0), especially when compared to the healthy controls.

### Classes of proteins displaying alike trends in plasma concentrations

To reveal possible relations between individual proteins, we present the quantitative proteomics data also in a protein-based dendrogram (**Figure 2b**) to extract which proteins cluster closely together, showing similar responses. The distance of the clusters is based on the complete linkage method, which calculates the distance of the farthest elements in the cluster. Proteins that reveal alike behaviour (complete distance less than 0.3) are depicted with the same colours and are separated by the red dashed line. Obviously, we observe all three chains of fibrinogen (FGA, FGB, FGG) coloured in light blue, tightly correlating, having a distance smaller than 0.1, indicating that they, as expected, form a well-defined stoichiometric protein complex, which makes their shifting concentrations very similar to each other. A similar tight cluster is seen for Complement C1q subcomponent subunits A, B and C (C1QA, C1QB, C1QC), that together form part of the 18-component protein assembly C1Q, and Complement component C8 alpha, beta and gamma chains (C8A, C8B, C8G), that form together C8^17, 18^. Also, the major red blood cell contaminant proteins like Hemoglobin subunit alpha and beta (HBA1, HBB), Peroxiredoxin-2 (PRDX2) and Carbonic anhydrase 1 (CA1), do cluster tightly. Other examples are the tightly connected SAA1 and SAA2, as are S100A8 and S100A9. These four latter proteins furthermore cluster closely to CRP. The latter is routinely used clinically as biomarker in the diagnosis of infections, autoimmune diseases, and cancers.

### Correlation between plasma proteomes

Next, we constructed a sample-oriented correlation dendrogram, wherein the distances in between samples direct reflect their alikeness considering all the measured concentrations of the 200 most abundant unique proteins in the plasma proteome (**Figure 3**). This dendrogram nicely reiterates the closeness in the plasma proteomes of all six control plasma proteomes. Interestingly, also the plasma proteome of P2_T4 correlates well with the healthy donors, indicating that the plasma proteome of patient P2 may have converged to a “normal” proteome at the latest sample point, a year after the kidney transplant. Also, this dendrogram reveals the alikeness of the two outlier plasma proteomes, sampled from the CKD patients prior to the transplants. Other samples also reveal tight correlations, most clearly when focusing on distinct sampling points for P1. P1_T1 and P1_T8 cluster tightly, which we later show to be at the height of APR events in P1. Similarly, P1_T3 and P1_T9 cluster tightly, which we later show to be both at minimum of APR events.

**Figure 3.**
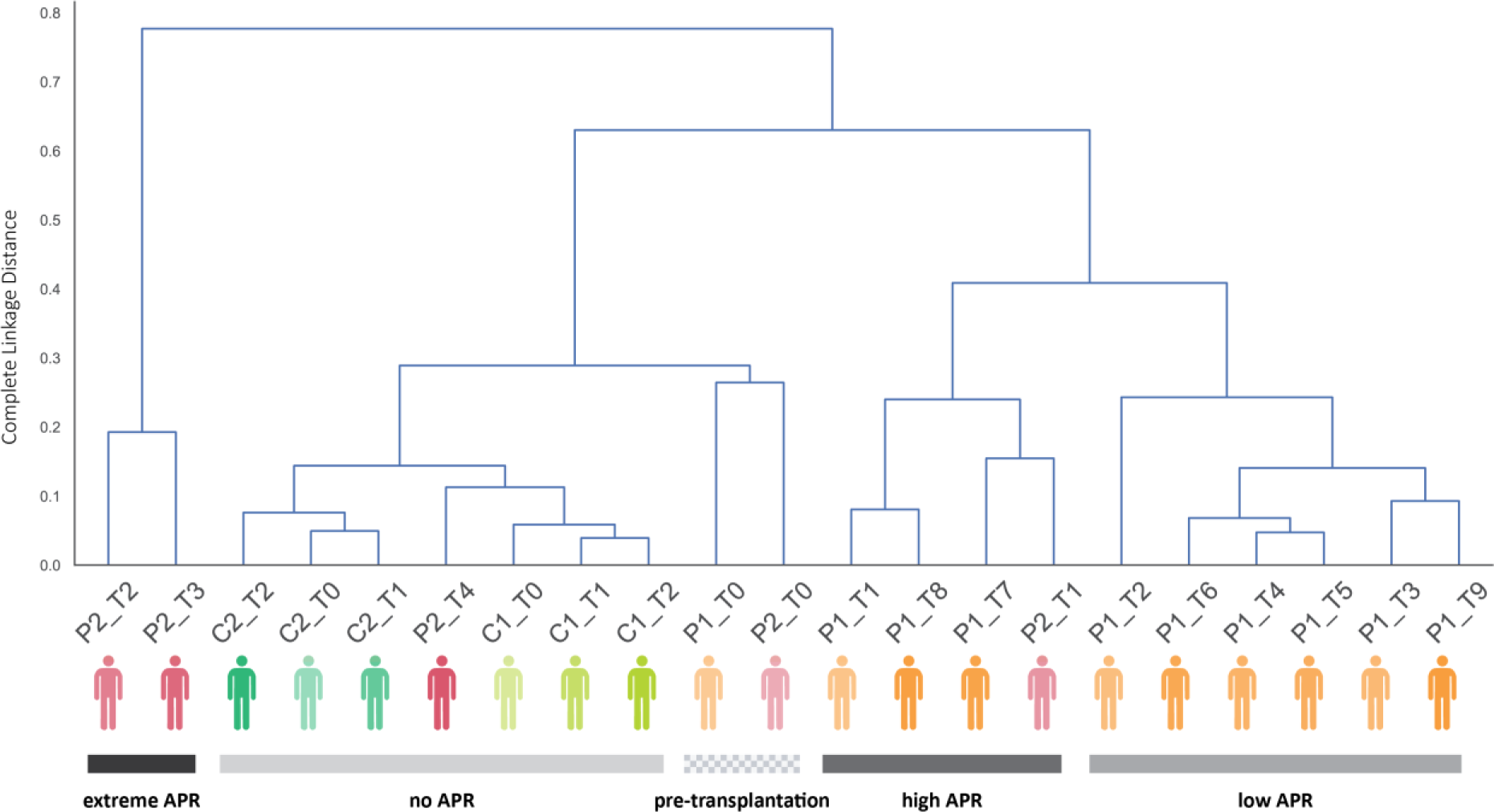
Clustering of different timepoints in Patients and Controls based on their proteome profile. In the dendrogram the individuals are coloured based on the colour scheme used in Figure 1 and we used an increasing opacity from T0 to the last timepoint (T0 faded, Tx intense). We also divided the dendrogram in five clusters, based on the phase of APR that the patients are in. The squared grey depicts the pre-transplantation timepoints where both patients have unique profiles but similar to each other. In the light grey cluster, we see all the controls and the T4 of P2, which indicates that the patients’ profile starts resembling the healthy.

Next, we looked at what made the plasma of the two CKD patients prior to the operation that distinctive and observed dramatic lower concentrations of the usually highly abundant proteins FGA, FGB, FGG (∼100 fold) and the immunoglobulins IGHG2, IGHG3, IGHG4, IGHA1 and IGHA2 (∼ 5-fold) in the patients prior to the operation. We speculate that this is possibly an effect of proteinuria, but also dialysis and other treatments given to the CKD patients may have affected that plasma proteome^19^. Notably, other abundant plasma proteins showed no substantial alterations between these two samples and the controls, including albumin, Serotransferrin (TF), Transthyretin (TTR), Hemopexin (HPX), Haptoglobin (HP), Histidine-rich glycoprotein (HRG), and Alpha-1-antitrypsin (SERPINA1).

### Plasma Complexome profiling by SEC-LC MS

Complexome profiling is a combination of size-exclusion chromatography (SEC) and LC-MS^12,20^. In our experiments we fractionated the plasma samples by SEC, whereafter consecutively all ∼ 60 fractions are analysed by bottom-up proteomics. In this way, in SEC co-eluting proteins can be identified, possibly leading to the identification of new interaction partners in protein assemblies. The molecular weight (MW) span of our approach allows to monitor proteins and assemblies in the mass range between 10 kDa and 3000 kDa. We performed complexome profiling on plasma samples from the controls C1 and C2 (T0, T1, T2) and from P2 (T0, T1, T2, T3, T4). The resulting SEC chromatograms are depicted in **Figure 4**. Some of the most abundant peaks in the chromatogram (annotated and centred around F20, F43-F47 and F55) are obviously linked to the most abundant proteins in plasma and were found by bottom-up proteomics to correspond predominantly to FG (F20-F22), A2M (F26-F28), HP (F45-F47), IGHGs/IGHAs (F53) and ALB (F60), respectively. While the SEC profiles were relatively alike in the healthy controls C1 and C2, in patient P2, striking changes were observed over time, with especially the signals in fractions F20-22 and F31-32, being much more abundant at T2 and T3. Detailed information about the SEC elution profiles of each of 135 individual plasma proteins is provided in Supplementary Table 2, a selection of which will be discussed below.

**Figure 4.**
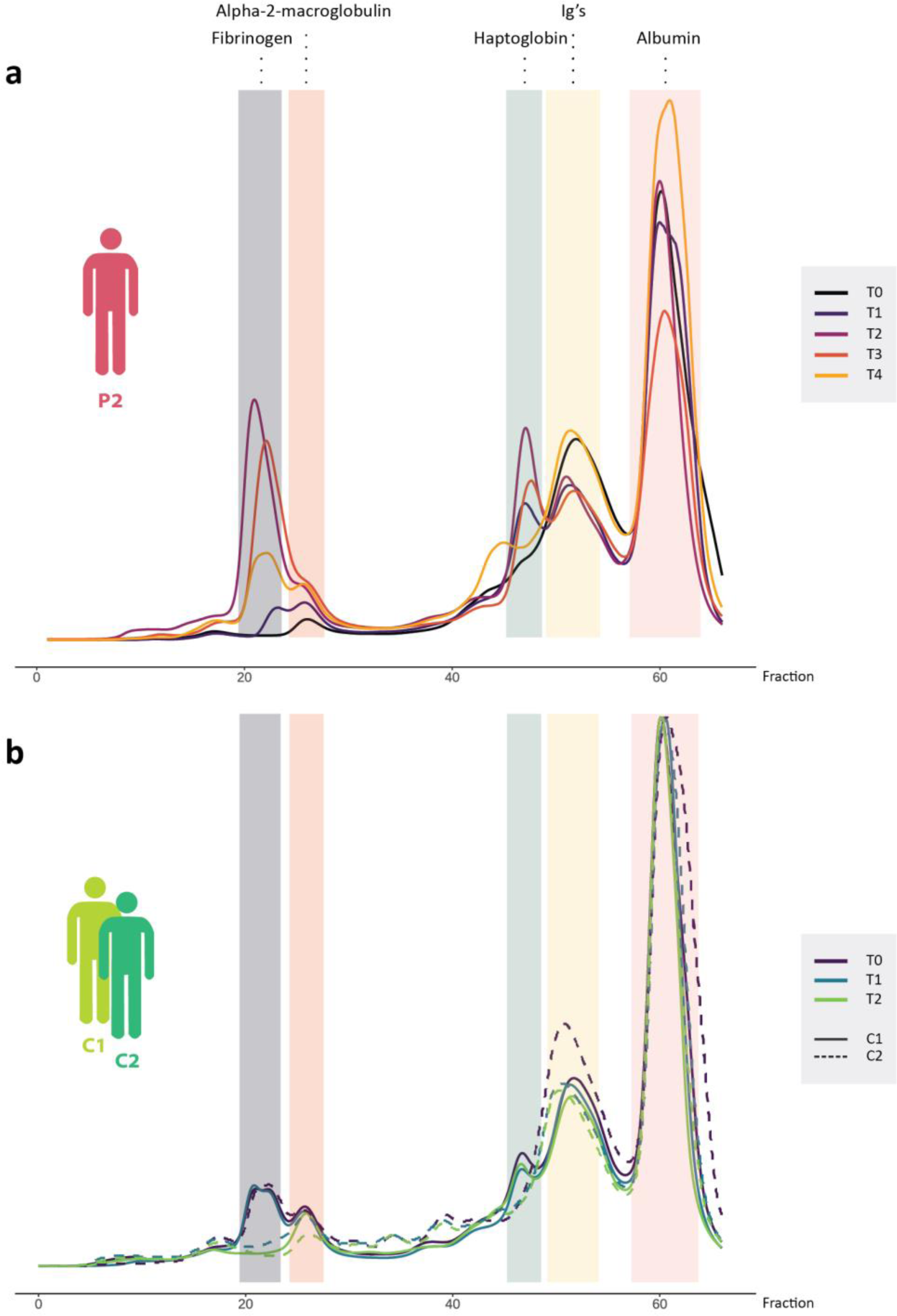
SEC chromatograms of plasma from C1, C2 and the P2 patient. The most intense peaks are highlighted and annotated by the most abundant proteins found in these fractions based on proteomics analysis. For example, the most intense peak in pink belongs to albumin and in the yellow fraction mostly Ig proteins are found. **a.** In the SEC elution profile at different sampling timepoints for P2 patient we observe substantial increases in FG and HP during APR. For the samples of the healthy individuals in **b.** we observe an overall stable elution pattern.

## Discussion

### Hallmarks of inflammation: positive and negative APPs

In the following discussion we zoom into proteins for which we observed interesting patterns in variation of their concentrations over time, especially in P1 and P2. As these CKD patients have suffered from bacterial infections, we first focused on hallmarks of inflammation.

CRP and SAAs are the best-known protein biomarkers for monitoring inflammation and acute phase responses^21, 22^. Human CRP is the most classical acute phase reactant, the concentration of which in plasma rises rapidly and extensively in a cytokine-mediated response to tissue injury, infection and/or inflammation^23^. Both CRP and SAA proteins are typically low abundant in plasma of healthy donors but rise 10-100 times in plasma concentration following inflammation. Consequently, we used these proteins (CRP, SAA1 and SAA2) to evaluate the inflammation along the time course of sampling. Indeed, we did find these three proteins to be low abundant and constant in concentration in the plasma samples from the healthy donors at all 3 time points with concentrations of around 0.3, 0.7 and 0.05 mg/dL for CRP, SAA1 and SAA2, respectively (**Figure 5a** and **5b**). Such alike low concentrations are also observed for the two patients at T0 (before the transplantations and infections) and at the latest sampling points (T10 for P1 and T4 for P2). Monitoring the concentration of these proteins over time in the patients reveals that P2 has especially at T2 highly elevated levels. Whereby, CRP concentrations have gone up 50-fold and SAA1 and SAA2 more than 100-fold, to concentrations of 50, 300, and 100 mg/dL for CRP, SAA1 and SAA2, respectively. For P1 the levels of these proteins also rise substantially, although less than in P2. For P1, we seem to observe multiple maxima in their concentrations, notably at T1 and T8. The maxima observed in the concentrations clearly reveal that the patient at time of these samplings experienced an APR, likely linked to the observed bacterial infections.

**Figure 5.**
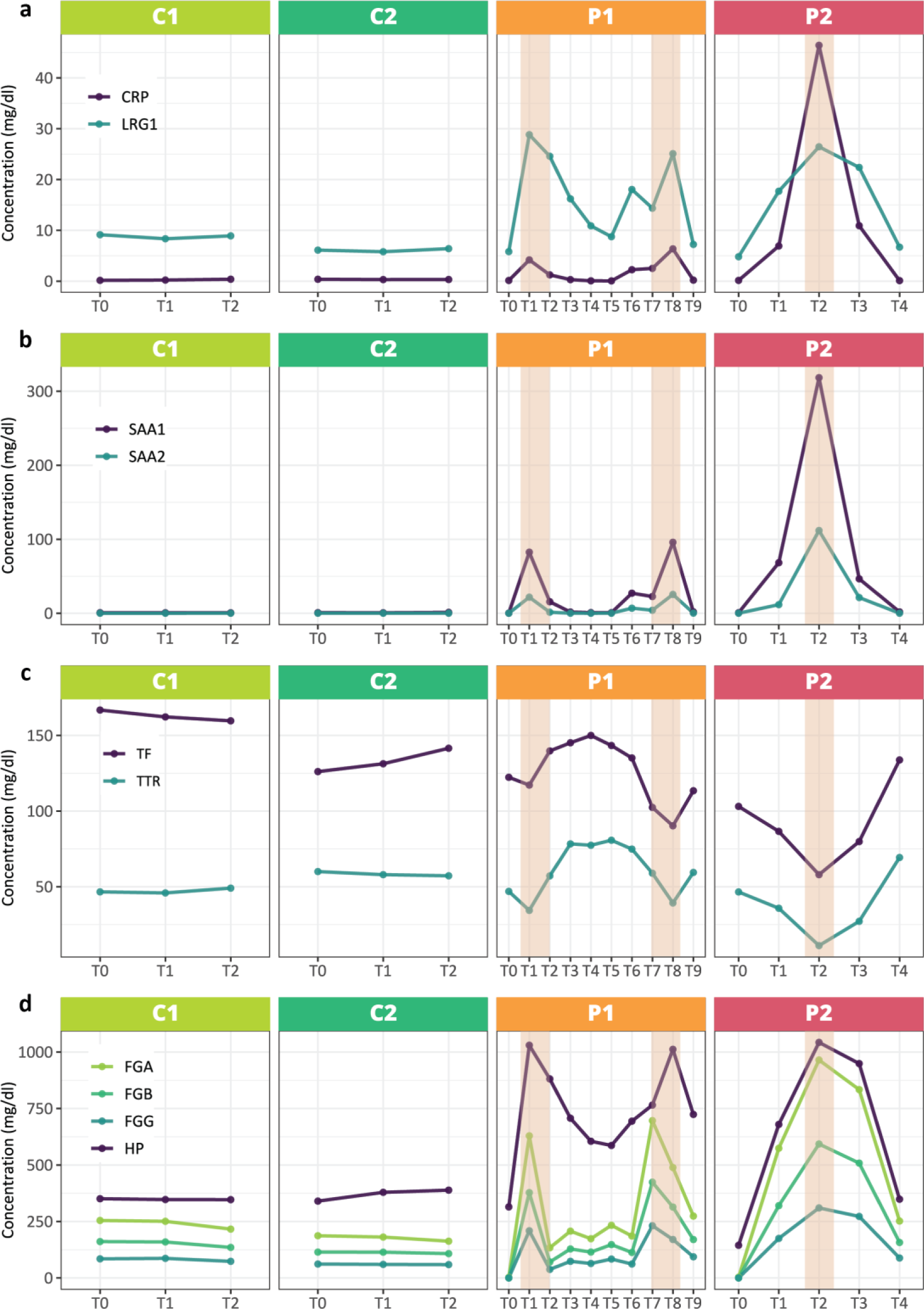
Plasma concentrations of selected positive and negative acute phase markers at different sampling timepoints for each donor. The regions highlighted in orange in P1 and P2 indicate the occurrence of bacterial infections. The healthy individuals show relatively stable profiles for all depicted proteins, whereas in the patients during the time of infection there are substantial variations observed.

CRP and SAA are well-known positive acute-phase proteins, whereas TF and TTR represent hallmark negative acute phase proteins. In our data the trends in concentrations of these proteins nicely display “mirror images” of the profiles of CRP, SAA1 and SAA2, with in P1 two minima at T1 and T8, and in P2 a strong minimum at T2 (**Figure 5c**). It should be noted that these negative APPs are also abundant in the healthy donors, and thus the observed minima represent decreases of just about 2-fold. This makes them less quantifiable/distinguishable biomarkers for inflammation than the above-described positive APPs, CRP and SAA. The SEC-LC-MS profiles of CRP (see **Supplementary Figure 2**) indicate that this small protein (MW ∼ 24 kDa) eluted at a MW of ∼ 125 kDa, indicative of its known presence as pentamers in plasma^23^. SAA1 and SAA2 represent even smaller proteins (MW ∼ 12 kDa) but it has been reported that native SAA1 exists primarily as a ∼ 70 kDa hexamer. The SEC-LC-MS profiles of SAA1 and SAA2 show that these two isoforms perfectly co-elute with each other, albeit in fractions that are more indicative of assemblies of in between 400-1000 kDa, observations to which we come back later. The data on these five hall-mark proteins unambiguously reveal that both patients experienced one or more APRs. Subsequently, we analysed the longitudinal concentration profiles of all detected plasma proteins, to see whether other proteins followed patterns alike these hallmark acute phase proteins.

### Additional (presumed) positive acute phase proteins Calprotectin; S100A8 and S100A9

The S100 proteins S100A8 and S100A9 are barely detectable in the plasma of the healthy controls with concentrations around 0.02 to 0.1 mg/dL. S100A8 and S100A9 form together the calprotectin hetero-dimer (MW ∼ 24 kDa dimer)^24^. Both subunits are rather small (MW ∼ 11 and 13 kDa) and structurally homologues. Calprotectin is regulating several inflammatory processes and the immune response. Therefore, it may come as no surprise that the plasma concentrations of both these proteins follow similar patterns as CRP and SAA in our data and rise in P1 about 10-fold and in P2 even up to 50-fold, with in P2 concentrations of S100A8 peaking around 20 mg/dL at T2 (**Supplemental Figure 1a**). Our SEC-LC-MS profiles indicate that both these proteins indeed co-elute and are present in plasma as calprotectin (S100A8/S100A9 hetero-dimers), although a small part seems to form calprotectin tetramers (i.e. hetero-tetramers), in line with what has been previously reported^24^. Since calprotectin levels in healthy donors are very low, these proteins may also be considered as decent biomarkers for inflammation.

### Leucine-rich α-2 glycoprotein 1 tightly co-varies with CRP

The also by hepatocytes secreted leucine-rich α-2 glycoprotein 1 (LRG1) has been implicated in multiple diseases, including cancer, diabetes, cardiovascular disease, neurological disease, and inflammatory disorders^25^. It has been postulated as an APP with rapidly increasing plasma concentrations following microbial infections and other inflammatory stimuli, although its APP status is not as recognized as CRP and SAA^26, 27^. In our data the temporal concentration profiles of LRG1 matched extremely well that of CRP (**Figure 5a**). Both CRP and LRG1 concentrations levels in plasma peaked around 50 mg/dL, at T8 for P1 and T2 for P2. The major difference between CRP and LRG1 in our data is that “baseline levels” in the healthy donors are around 0.3 mg/dL for CRP and around 6-8 mg/dL for LRG1. Therefore, although the absolute rise in abundance upon inflammation of CRP and LRG1 is quite similar, the fold changes are quite different. LRG1 was found to elute late in the SEC-LC-MS profiles, likely just as monomer (Supplementary Table 2).

### Fibrinogen and Haptoglobin

FG and HP are among the highest abundant proteins in plasma with, also according to our data, concentrations of ∼ 100’s of mg/dL. Fibrinogen consists in plasma of three subunits, FGA, FGB and FGC, forming together predominantly 2:2:2 hexamers of around 375 kDa^28^, whereas in plasma HP (depending on the allotype) can form homo- and hetero-oligomers that may bind strongly to HB ^6^. The plasma concentrations of all these proteins do reveal very similar patterns in our data, that also to a large extend mimic those observed for CRP and SAA (**Figure 5d**). Overall, it seems that fibrinogen concentrations rise 3-to-5-fold at maxima of inflammation (T1 and T8 in P2, and T2 in P2), whereas HP concentrations have gone up 2- 3-fold at these same maxima, coming back to levels as observed in the healthy controls at the other timepoints. At these maxima of inflammation, the levels of fibrinogen and HP reach a concentration close to what we measure for albumin, making them also noticeably observable in the SEC chromatograms displayed in **Figure 4**. Fibrinogen and HP therefore seem to follow in the CKD patients the hallmark of positive acute phase proteins, such as CRP and SAA, but are likely less distinctive biomarkers as they are already highly abundant under normal physiological conditions. The SEC-LC-MS profiles indicate that as expected the three chains of FG (FGA, FGB, FGG) co-elute quite early in the SEC profile, indicative of being present in its known hetero-hexameric A2B2G2 form^28^. The elution profile of HP in the SEC-LC-MS data of P2 was around 150 kDa, consistent with the mass of a HP1-dimer. Indeed, we extracted from the proteomics data that P2 is homozygote for HP1-1.

### The protease inhibitors alpha-1-antitrypsin and alpha-1-anti chymotrypsin

Alpha-1-antitrypsin (A1AT, or SERPINA1) and alpha-1-anti chymotrypsin (AACT, or SERPINA3) represent some of the most abundant serpins in plasma, acting primarily as protease inhibitors. In the healthy controls C1 and C2 the levels of these two protease inhibitors remain longitudinally constant at levels of about 200 μg/dL and 25 μg/dL, for A1AT and AACT, respectively. In the patients we observe patterns alike those of other positive acute phase proteins (**Supplementary Figure 1b**). At the maxima the levels of A1At and AACT rise about 2-to-5 fold, to about 500 μg/dL and 120 μg/dL, for A1AT and AACT, respectively. These findings corroborate A1AT and AACT as being positive acute phase proteins. These two proteins are often proposed as biomarkers in plasma proteomics studies, not only when studying bacterial infections, but also when distinguishing healthy donors from cancer patients^29–33^. Our SEC-LC-MS profiles indicate that both these proteins are largely present in plasma as monomers although a small part of plasma A1AT (less than 5%) co-eluted in higher MW fractions (Supplementary Table 2).

### LPS-binding protein (LBP) and its ligand CD14

LPS-binding protein (LBP) with its ligand CD14 are located upstream of the signalling pathway for LPS-induced inflammation. Preventing LBP and CD14 from binding supposedly hinders LPS-induced inflammation^34^. Both these proteins are relatively low abundant in plasma in healthy donors with concentrations of about 0.6 and 0.2 mg/dL for LBP and CD14, respectively. In our proteomics data for the patients P1 and P2 we observed substantial higher levels at the same maxima as seen for the hallmark positive acute phase proteins, the patterns matched extremely well that of e.g., CRP and LRG1 (**Supplementary Figure 1c**). At the maxima of inflammation levels of these proteins went up about 3-to-8 fold, with a more sizeable increase for LBP compared to CD14. Therefore, we propose that these proteins may also be regarded as positive acute phase proteins, albeit that they are orders less abundant in plasma than some of the positive acute phase proteins described above.

### Additional (presumed) negative acute phase proteins

Based on their concentration profiles we propose that inter-alpha-trypsin inhibitors (I*α*I), Heparin cofactor 2 (SERPIND1) and Kallistatin (SERPINA4) can be regarded as putative negative acute phase proteins (**Supplementary Figure 1c**). Their concentration profiles follow the trends observed for TF and TTR, in line with previous reports indicating that I*α*I plasma concentrations decline during acute inflammation^35^. Notably, IαI has been used for replacement therapy in the treatment of patients upon inflammatory conditions. The concentration profiles of the ITIH1 and ITIH2 correlate very well (**Figure 1** and **Supplementary Figure 1b**), which can be explained as they are known to form in plasma an approximately 225-kDa complex, named IαI, containing Bikunin (C-terminal fragment of Alpha-microglobulin/bikunin precursor (AMBP)) next to ITIH1 and ITIH2^36^. In our SEC-LC-MS complexome data we indeed observed also the perfect co-elution of ITIH1 and ITIH2 in a fraction that could well represent a ∼ 225 kDa complex (**Supplementary Figure 3**). The third subunit AMBP also co-eluted with ITIH1 and ITIH2 but is also present in other fractions, likely as it can also bind to other proteins (e.g., ITIH3).

Two other serpins clearly showed patterns alike negative acute phase proteins, namely SERPINA4 and SERPIND1. SERPINA4 inhibits amidolytic and kininogenase activities of tissue kallikrein, and has anti-peptidase activity, but not much more is known about this protein. SERPIND1, is also known as Heparin cofactor 2 and is an important thrombin inhibitor.

### Observation of dynamic changes in HDL particles during inflammation

In plasma there are various particles consisting of lipids and proteins whose function it is to transport lipids throughout the body in blood^37^. These lipoprotein particles typically consist of a core containing cholesterol esters and triglycerides surrounded by free cholesterol, phospholipids, and apo-lipoproteins^38^. Here, we focus on the high-density lipoprotein (HDL) particles for which the protein composition is dominated by its structural apolipoproteins APOA1, APOA2, and APOA4 (**Figure 6a**)^39^. However, HDL particles are quite dynamic in composition as both lipids as well as proteins can be interchanged, depending on the physiological condition of the donor^40^. In our quantitative proteomics data, we observed trends for these APOA proteins that are very much alike those of the negative acute phase proteins, described above, with clear minima in protein concentrations at T1 and T8 in P1 and T2 for P2 (**Figure 6b**). Additionally, we plotted in **Figure 6b** the protein concentration as measured for SAA1 and SAA2, and it may be hypothesized that the loss in concentrations observed for the APOA proteins, at the height of APR, could be compensated by the gain in concentrations of SAA1 and SAA2. This hypothesis is not completely novel, as it has been reported that during a systemic inflammatory response, SAA can be incorporated into high-density lipoprotein (HDL), and even become the major apolipoprotein on HDL^41, 42^. Our data support this finding explicitly. Next, we also evaluated the co-elution profiles of these proteins in our complexome data and observed indeed that SAA1 and SAA2, that when purified form just monomers or small oligomers^40^, elute in the SEC-LC-MS data in high molecular weight fractions (MW ∼ 250 kDa), largely overlapping with the fractions in which also the APOA1, APOA2 and APOA4 proteins are observed (**Figure 6c**).

**Figure 6.**
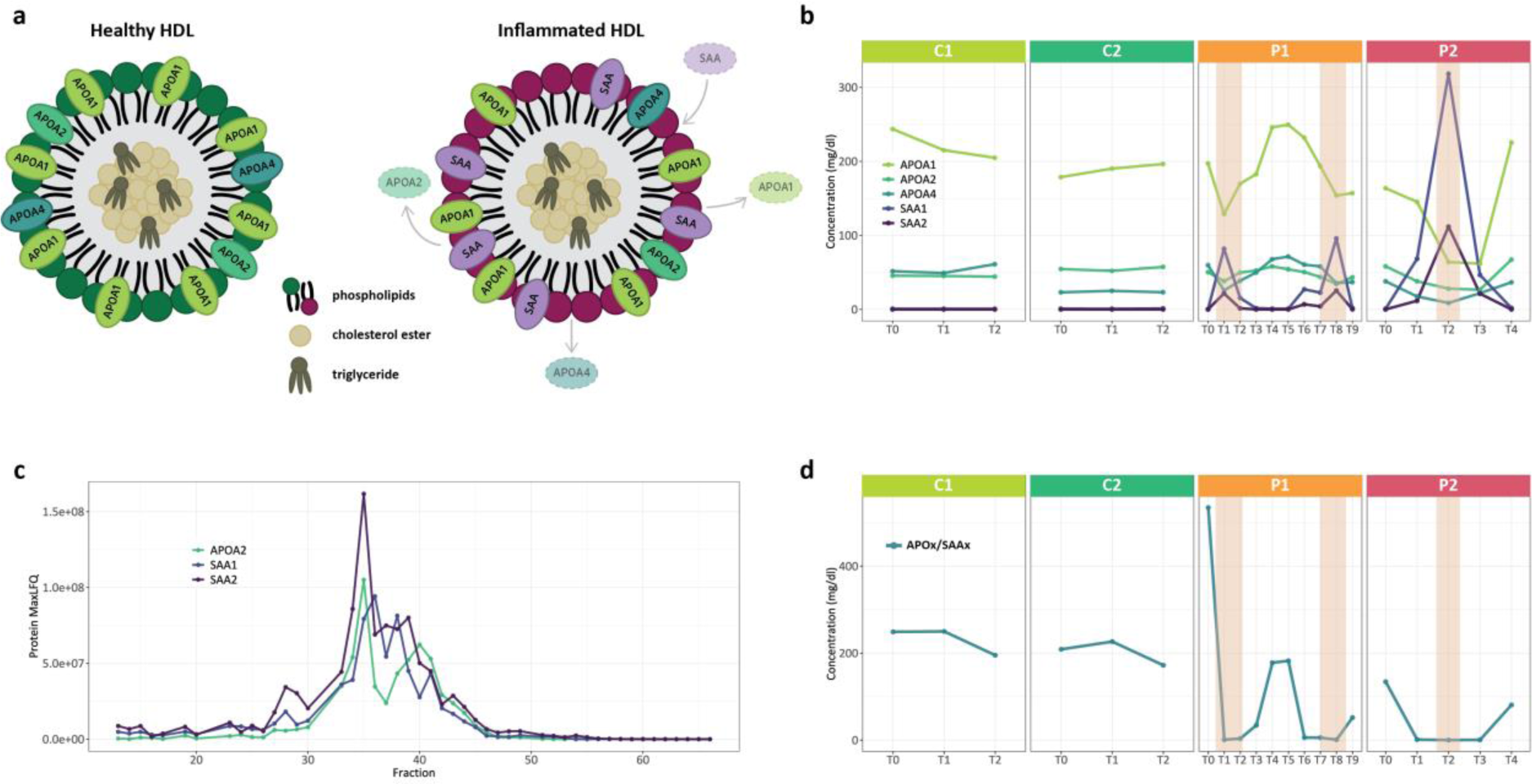
Dynamic changes of apolipoproteins and serum amyloid proteins in plasma HDL particles during APR. a. Model of the composition of HDL particles under normal and inflamed conditions. During APR APOx proteins may be replaced by SAAx. This model is confirmed by our data in **b.** where we observe the concentrations of APOx and SAAx in the controls to be stable while in the patients there is a decrease of APOx and concomitant increase of SAAx at the APR maxima. **c.** The elution profiles from the SEC data of the P2 patient at the time of inflammation (T2) reveal the coelution of APOA2, SAA1 and SAA2 in HDL particles. **d.** Representation of the concentration ratio of the sum of APOA1, APOA2, APOA4 divided by the sum of SAA1 and SAA2, which may be considered an even more distinctive marker for APR. The regions highlighted in orange in P1 and P2 indicate the occurrence of bacterial infections.

Our study indicates that also APOA1, APOA2 and APOA4 could be considered negative acute phase proteins, but again as they are already high in concentration under normal physiological conditions a decrease in in abundance may be harder to measure. Evidently, when one could assess the ratio APOAx/SAA (as depicted in **Figure 6d**), this could be regarded as a superior and more sensitive biomarker for inflammation. Moreover, our data reveal a somewhat equal propensity for replacement of APOA1, APOA2 and APOA4 by SAA1/SAA2 in HDL particles under conditions of acute phase.

## Summary of Strengths and Weaknesses

Here we report an in-depth quantitative plasma proteomics study, using an alternative approach when compared to several recent high-throughput plasma proteomics studies that use the power of proteomics to study the plasma proteome in large cohorts of donors^1–4^. A major weakness of our study is that it is limited to just four donors, two healthy donors (C1, C2) and two CKD patients (P1, P2). The latter two were the focus of this study, as they both endured a kidney transplant. Therefore, especially these patients were carefully monitored and revisited the hospital/doctors for numerous check-ups over a timespan of a year, enabling us to obtain longitudinal samples of their blood. Evidently, our study would be strengthened by including more patients, but also patients for which we had even more sampling timepoints. Unfortunately, such samples are not easy to obtain, unless a specific new cohort would be started. A strength of our longitudinal study is that we make a strong case for such future “bio- banking” initiatives that would sample blood at much more regular intervals for several patients and follow these up with quantitative plasma proteomics to monitor the effects of an operation, infection, therapeutic treatment etc., possibly enabling doctors to adjust treatments.

Most of the proteins that we observe to be changing are well-known acute phase response proteins, and therefore we report limited new biology. Still, we show that in-depth quantitative plasma proteomics has now matured so much that all such acute phase proteins can be robustly and quantitatively measured accurately in a multiplexed manner by proteomics. Moreover, we validate some lesser- known APR proteins as genuine members. Next to the well- known acute phase proteins (CRP, SAA, TF and TTR) this study provides further support for S100A8/S100A9, SERPINA1, SERPINA3, LBP and CD14 as positive APR proteins and ITIH1/ITIH1 and APOA1/APOA2/APOA4 as negative APR proteins.

As a note of caution, our analysis reconfirms that there are dozens of proteins in plasma that change abundance, caused primarily due to an acute phase response and/or inflammation. Knowing this list of APR proteins well, may help to distinguish in other biomarker studies, e.g., related to cancer, genuine cancer related protein biomarkers from those that change their abundance due to a parallel occurring APR.

For the controls C1 and C2 and patient P2, we performed further in-depth analysis of the plasma proteome by complexome profiling using SEC to fractionate the plasma into 66 fractions that were all analysed by bottom-up DIA based LC-MS. This allowed us to define (co)elution profiles of over 165 plasma proteins and led to several interesting observations. Most notably, in P2 at the height of inflammation the small positive acute phase proteins SAA1 and SAA2 co-eluted in a SEC fraction that corresponds to over 250 kDa (whereas their MW is just ∼ 12 kDa). We rationalized this behaviour, by postulating that they become incorporated into HDL particles, partly and largely replacing the dominant structural HDL proteins APOA1, APOA2 and APOA4. Indeed, these three APOA proteins were found to largely coelute with the SAA1/SAA2 proteins in SEC.

In general, by focusing here on a small number of patients, albeit by monitoring in-depth the concentration levels and SEC elution profiles of hundreds of plasma proteins longitudinally, we present an alternative workflow for plasma proteomics that may help to identify better and more specialized protein biomarkers. Our study makes a strong case for more extensive longitudinal proteomics studies, that should not only focus on larger cohorts, but also on as many timepoints of sampling as possible, to benchmark the plasma proteome of each diseased donor, ideally against itself, when the donor was still (or again) healthy.

## Data availability

The raw LC-MS/MS files and analyses have been deposited to the ProteomeXchange Consortium via the PRIDE partner repository with the dataset identifier PXD046550. The R script for the conversion to concentrations is publicly available and can be accessed through this link https://github.com/hecklab/kidney_plasma.

## Acknowledgements

This research was partly funded by the Dutch Research Council through the NWO TTW-NACTAR Grant #16442 (to A.J.R.H. and S.H.M.R.). We additionally acknowledge support from NWO, funding the MS facilities through the X-omics Road Map program (project 184.034.019).

We acknowledge Andris Jankevics for his suggestions in the statistical analysis and Tim S. Veth for the help and advice with the instrumentation and the data processing. Tereza Kadavá and Shelley Jager are acknowledged for critically evaluating the manuscript.

## Compliance with ethical standards

All participants provided written informed consent to collect clinical data and blood samples. The study was approved by the local Biobank Research Ethics Committee (protocol 15-019) of the Utrecht UMCU.

## Supplementary figures

**Supplementary Figure 1.**
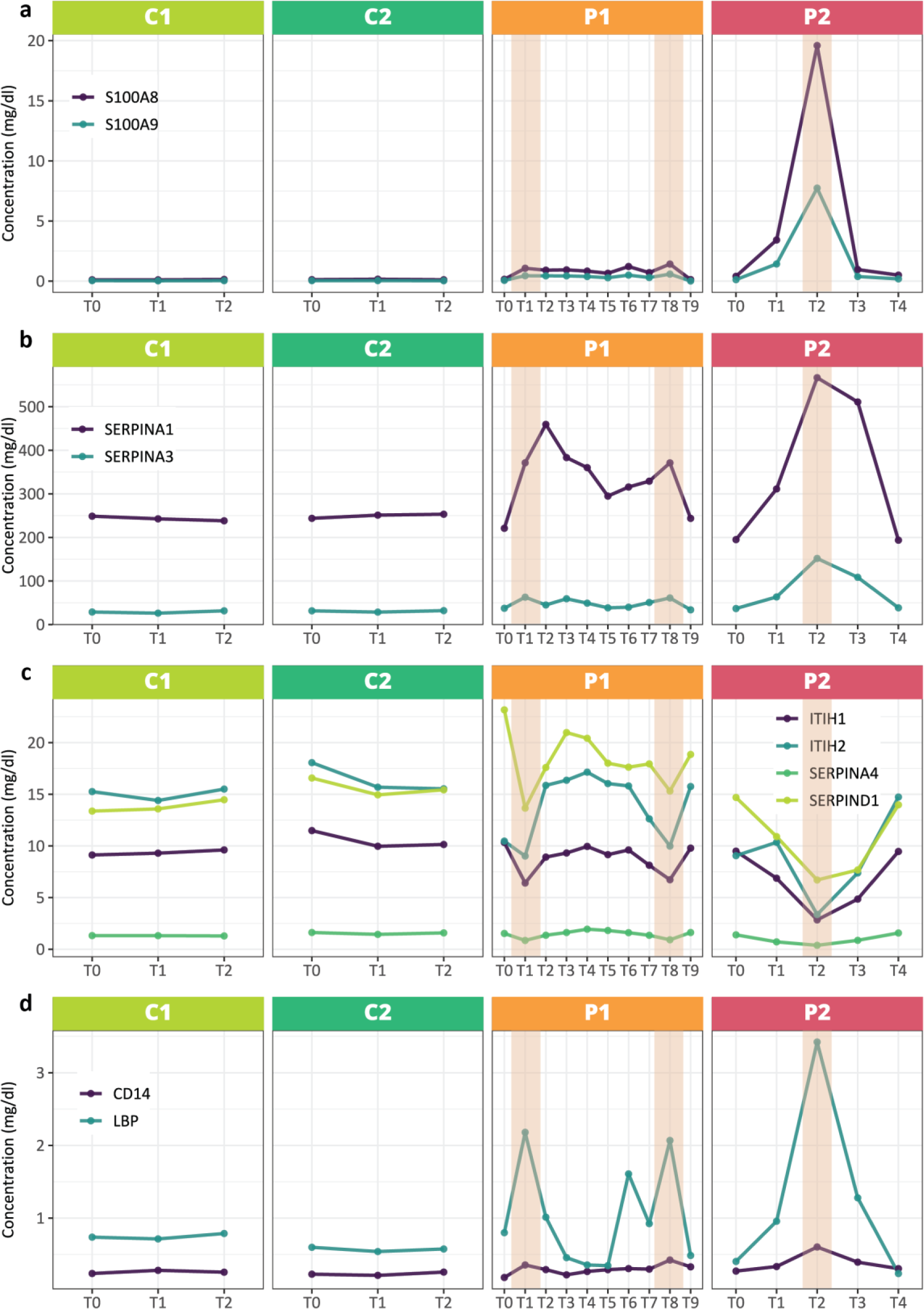
Plasma concentrations of selected proteins at different timepoints of sampling for each donor. The regions highlighted in orange in P1 and P2 indicate the occurrence of bacterial infections. The healthy individuals show relatively stable profiles for all depicted proteins, whereas in the patients during the time of infection there are substantial variations observed.

**Supplementary Figure 2.**
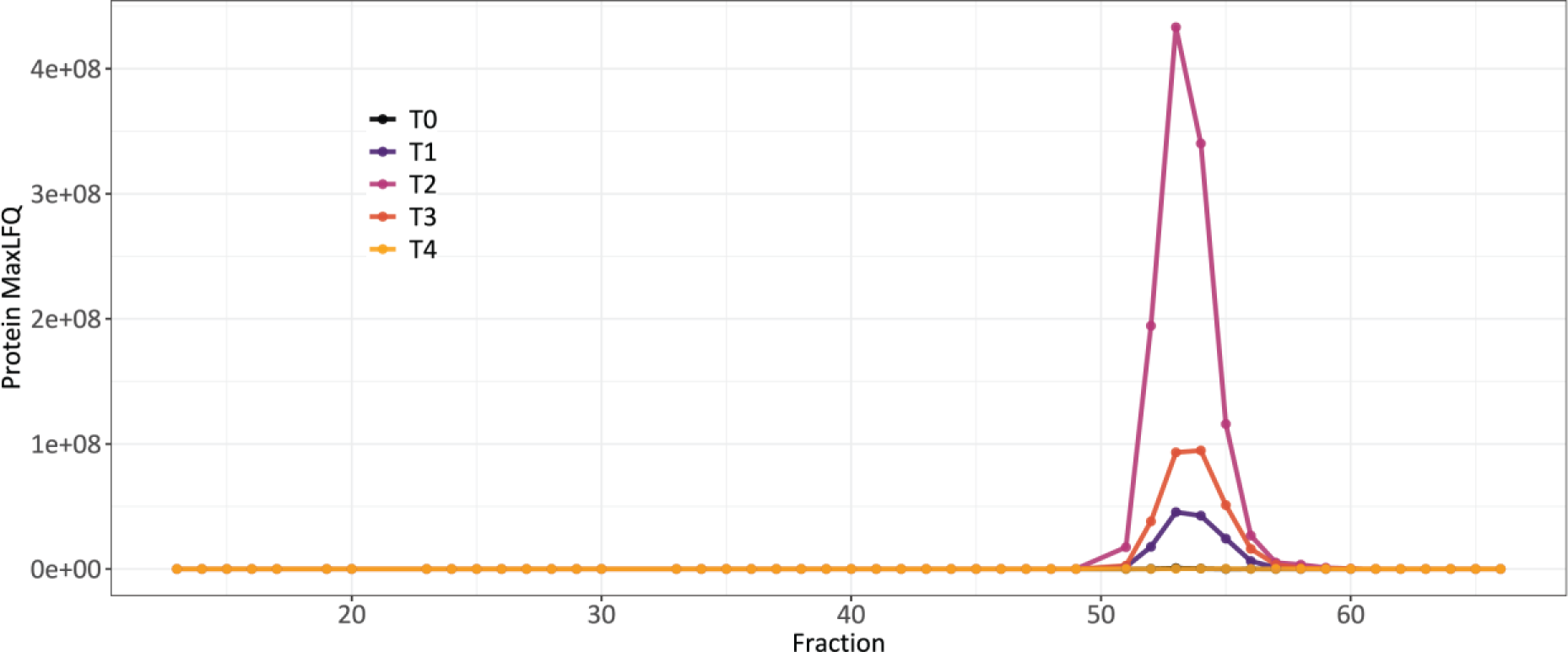
SEC LC-MS elution profile of CRP at distinct timepoint of sampling in P2. CRP elutes around fraction 55, consistent with the mostly pentameric form of CRP known to be present in plasma of ∼ 125 kDa.

**Supplementary Figure 3.**
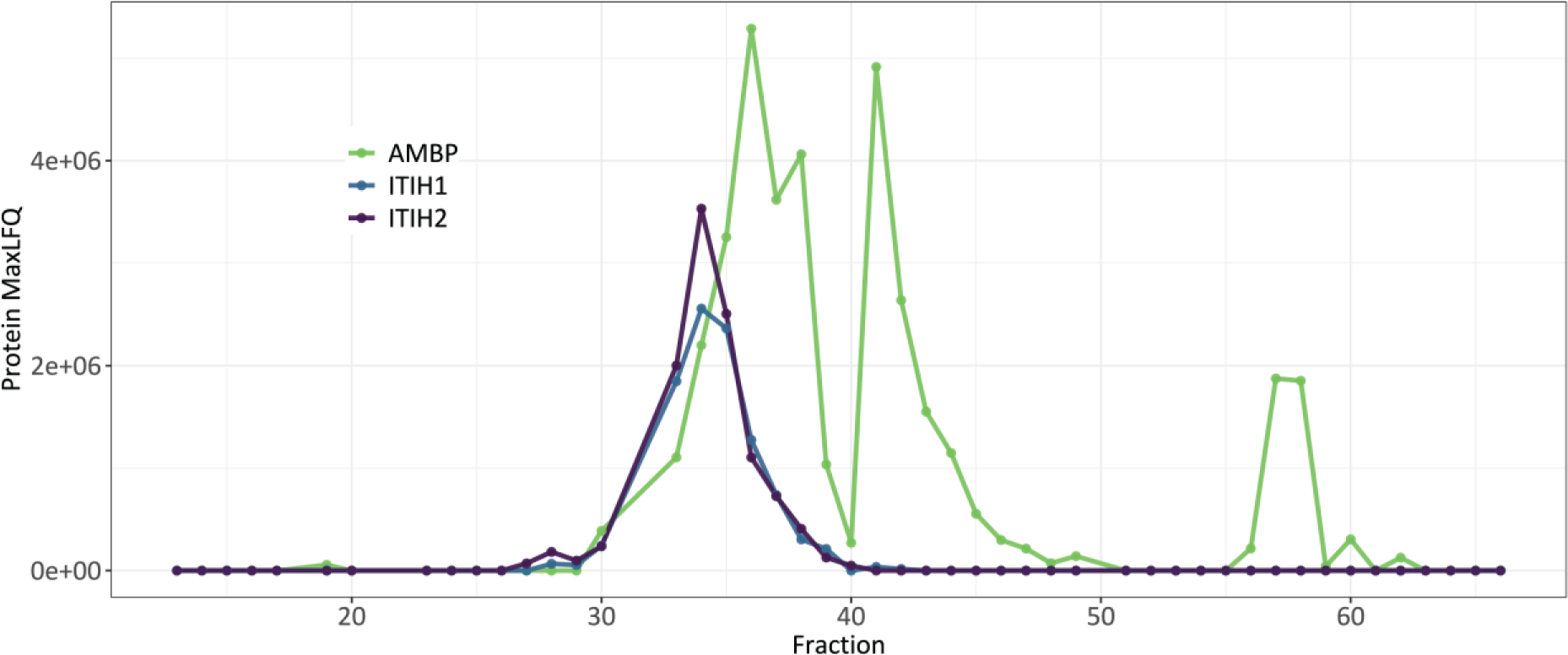
SEC LC-MS elution profile of ITIH1, ITIH2 and AMBP. ITIH1 and ITIH2 perfectly co-elute in fractions between 30 and 40, partially overlapping with the elution profile of AMBP. These three proteins are known to form together a complex in plasma of approximately 225-kDa complex, named IαI, containing the C-terminal fragment of AMBP next to ITIH1 and ITIH2.

