## Supplementary figures and images for "Longitudinal fluctuations in protein concentrations and higher-order structures in the plasma proteome of kidney failure patients subjected to a kidney transplant"

### supplemental Figure 1

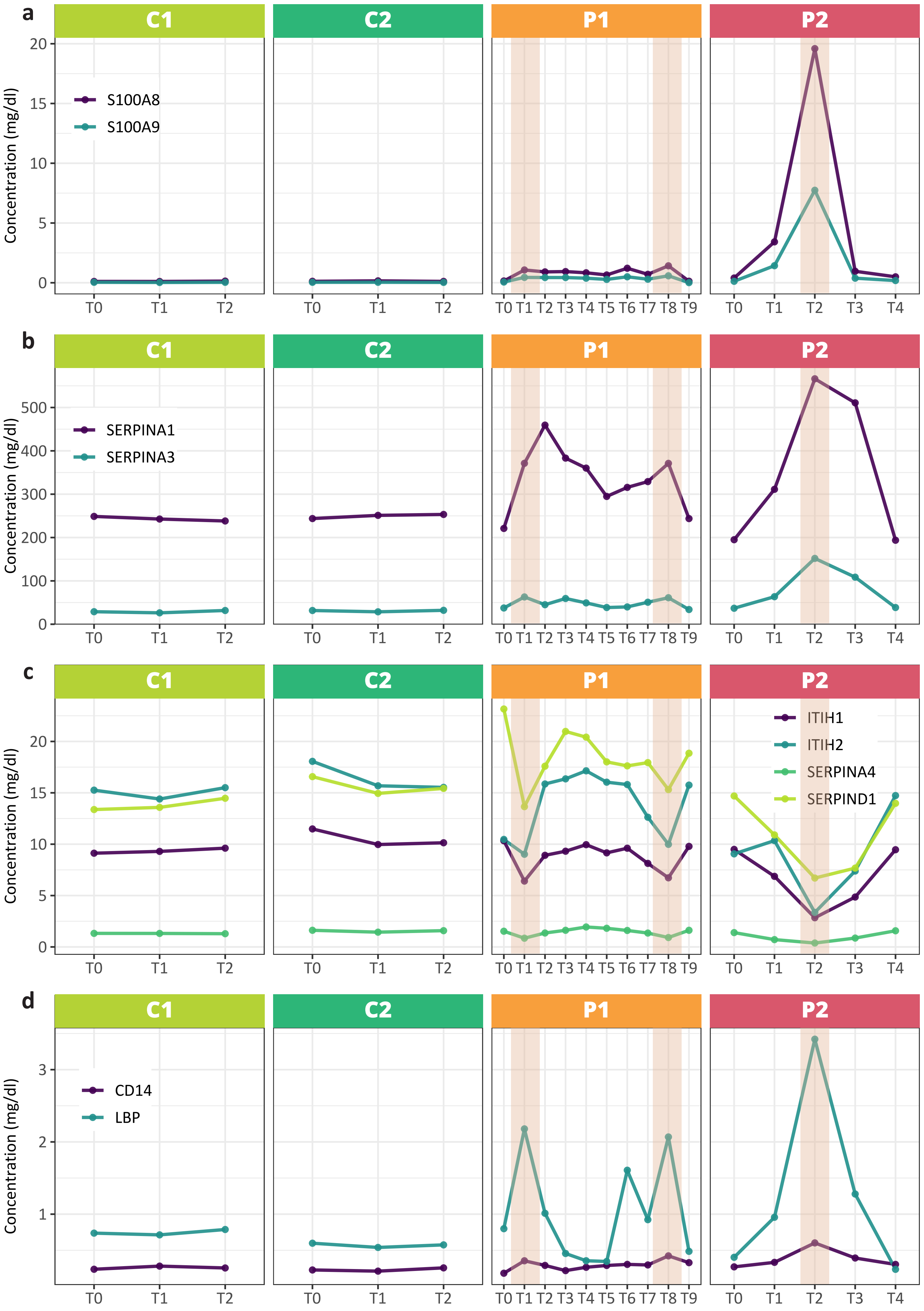

### supplemental Figure 2

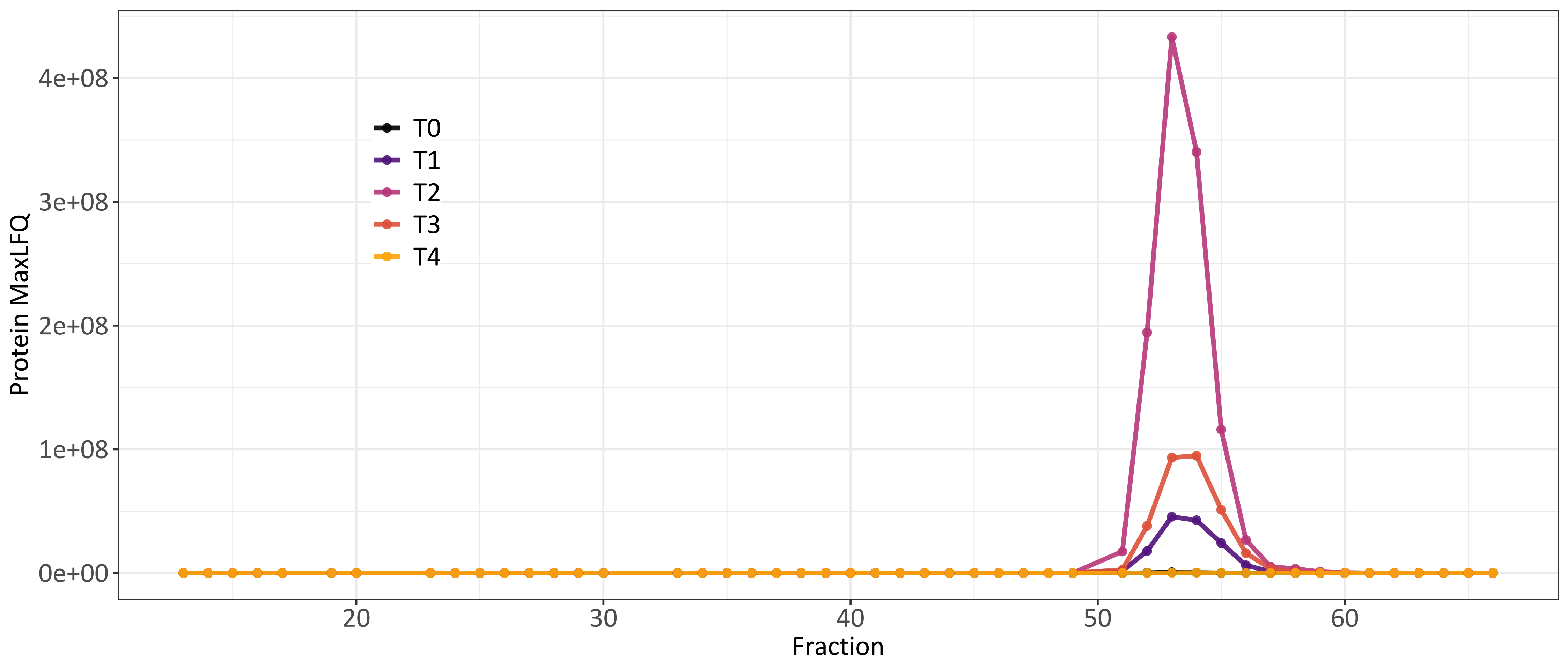

### supplemental Figure 3

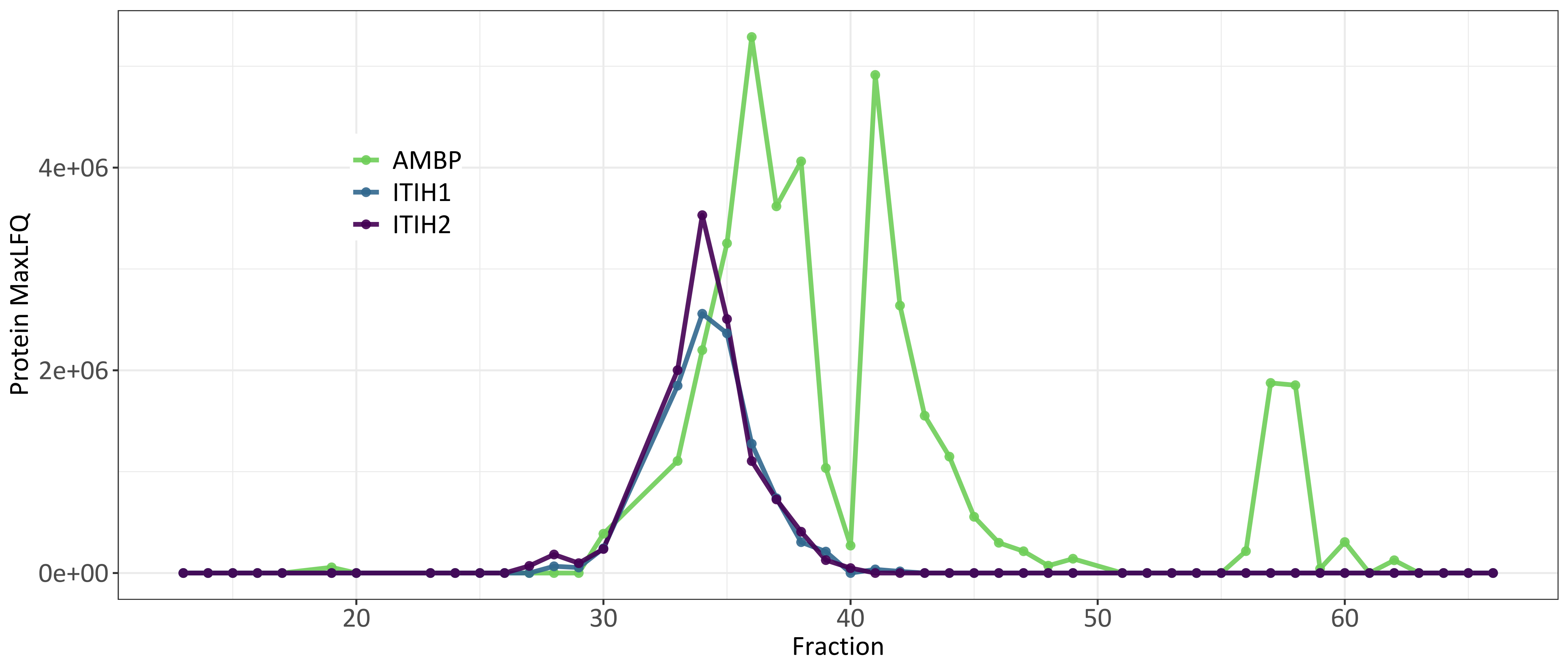
